# Demography, genetic, and extinction process in a spatially structured population of lekking bird

**DOI:** 10.1101/705301

**Authors:** Hugo Cayuela, Jérôme G. Prunier, Martin Laporte, Jérôme Gippet, Laurent Boualit, Françoise Preiss, Alain Laurent, Francesco Foletti, Gwenaël Jacob

## Abstract

Understanding the mechanisms underlying biological extinctions is a critical challenge for conservation biologists. Both deterministic (e.g. habitat loss, fragmentation) and stochastic (i.e. demographic stochasticity, Allee effect) demographic processes are involved in population decline. Simultaneously, a decrease of population size has far-reaching consequences for genetics of populations by increasing the risk of inbreeding and the effects of genetic drift, which together inevitably results in a loss of genetic diversity and a reduced effective population size (*N*_*e*_). These genetic factors may retroactively affect vital rates (a phenomenon coined ‘inbreeding depression’), and therefore reduce population growth and accelerate the extinction process of small populations. To date, few studies have simultaneously examined the demographic and genetic mechanisms driving the extinction of wild populations, and have most of the time neglected the spatial structure of populations. In this study, we examined demographic and genetic factors involved in the extinction process of a spatially structured population of a lekking bird, the western capercaillie (*Tetrao urogallus*). To address this issue, we collected capture-recapture and genetic data over a 6-years period in Vosges mountains, France. Our study showed that the population of *T*. *urogallus* experienced a severe decline between 2010 and 2015. We did not detect any Allee effect on survival and recruitment. By contrast, individuals of both sexes dispersed to avoid small leks, suggesting a behavioral response to a mate finding Allee effect. In parallel to this demographic decline, the population showed a low genetic diversity and high inbreeding. In addition, the effective population sizes at both lek and population levels was low. Despite this, we did not detected evidence of inbreeding depression: neither survival nor recruitment were affected by individual inbreeding level. Our study underlines the benefit from combining demographic and genetic approaches to investigate processes that are involved in biological extinctions.

## Introduction

In a global context of biodiversity decline, understanding the mechanisms underlying biological extinctions is a critical challenge for conservation biologists (Purvis 2000, Naeem et al. 2012, Pimm et al. 2014). Over the last three decades, many experimental and field studies investigated the demographic and genetic mechanisms involved in extinctions (reviewed in Young et al. 2000, Nunney & Campbell 1993, Frankham 2005). However, the relative contribution of these two types of mechanisms has been greatly debated for a long time and still remains controversial (Lande 1988, Nunney & Campbell 1993, Spielman et al. 2004, Frankham 2005). On the one hand, theoretical and empirical studies showed that populations are subject to extinction due to genetic factors, even in the absence of any human impact and ecological factor (Lande 1998, Szűcs et al. 2017). On the other hand, studies showed that demographic processes will generally doom small populations to extinction before genetic effects act strongly (Lande 1988, Wootton & Pfister 2013). Investigating the role of both demographic and genetic factors, and their interplay, thus remains an important challenge to better predict population extinctions and inform conservation strategies.

Both deterministic and stochastic demographic processes are involved in population decline (Lande 1993, Melbourne & Hastings 2008). Deterministic factors such as over-exploitation and habitat loss and degradation decrease population growth, which may ultimately lead to population extinctions (Selwood et al. 2015). Under stochastic processes, extinctions are caused by a high variance in population growth rate due to environmental stochasticity (e.g. weather fluctuations, catastrophic events) (Lande 1993, Lande & Orzack 1998, Engen et al. 2003) or demographic stochasticity (i.e. random variation of demographic rates) – whose effect increases when population size decreases. Moreover, extinctions may also be caused by density-dependent processes (Courchamp et al. 1999). In small populations, growth rate increases with density (i.e. positive density-dependence), a phenomenon coined the ‘Allee effect’ (Stephens et al. 1999, Stephens & Sutherland 1999). At low population density, individuals experience a fitness loss due the difficulty to locate mate (‘mate finding Allee effect’, Gascoigne et al. 2009) and/or to realize cooperative behaviors. This further translates into a decreased population growth and a higher risk of population extinction (Courchamp et al. 1999, Stephens et al. 1999).

Demographic processes described above have far-reaching consequences on population genetics (Tanaka 1997, Frankham 2005, Jaquiéry et al. 2009). A severe drop of population size is often associated with increased probability of breeding with relatives (i.e. inbreeding). Population decline also enhances the effects of genetic drift, which inevitably results in a loss of genetic diversity (Crnokrak & Roff 1999, Gibbs 2001, Frankham et al. 2002). Decreasing population size (*N*, often called ‘census population size’ in genetic studies) also results in a reduction in the effective population size (*N*_*e*_), a phenomenon that can be enhanced by the characteristics of the mating system (i.e. monogamy or highly skewed mating success) and several demographic features (e.g., overlapping generations or age structure) (Engen et al. 2005, Waples & Yokota 2006, Stubberud et al. 2017).

These genetic factors may retroactively affect vital rates (i.e. survival and recruitment; a phenomenon coined ‘inbreeding depression’), and therefore reduce population growth and accelerate the extinction process of small populations (Saccheri et al. 1998, O’Grady et al. 2006, Luque et al. 2016). Inbreeding may have a detrimental impact on adult survival (Meagher et al. 2000, Szűcs et al. 2017), fecundity (Westemeier et al. 1998) and offspring survival (Ballou & Ralls 1982, Keller 1998, Briskie & Mackintosh 2004, Spottiswoode & Møller 2004, O’Grady et al. 2006), and may be conducive to a loss of average fitness at the population level (Saccheri et al. 1998, Reed 2005). Simultaneously, a loss of genetic diversity, and therefore of quantitative trait heritability, may lead to a decrease of adaptive potential and possible population extinctions (Frankham 2005), even if the correlation between neutral genetic diversity and adaptive potential is usually low in natural populations (Reed & Frankham 2001).

To date, relatively few studies have simultaneously examined the demographic and genetic mechanisms driving the extinction of wild populations (Landguth et al. 2014, Mathieu-Bégné et al. 2018). Yet, those few studies have often neglected the spatial structure of populations. Most natural populations occur in heterogeneous, fragmented landscapes and are therefore spatially structured, i.e., composed of subpopulations occupying more or less discrete habitat patches linked together by dispersal (Thomas & Kunin 1999). Dispersal may play a critical role in population extinction by affecting both demographic and genetic processes (Thomas & Kunin 1999, Ronce 2007, Benton & Bowler 2012). At the individual level, dispersal ‘decisions’ depend on phenotypic traits (e.g. sex, age, social or breeding status), as well as social (i.e. population density, kin competition, inbreeding avoidance) and environmental factors (e.g. patch and landscape characteristics) (Clobert et al. 2009, Matthysen 2012). This results in non-random and generally asymmetric dispersal rates that mitigate the risk of subpopulation extinction (Hill et al. 2002, Gilpin 2012). At the demographic level, dispersal reduces the risk of subpopulation extinction through immigrant inflow (i.e. ‘rescue effect’, Hanski et al. 1997); immigration increases the size of a subpopulation and thus reduces Allee effect and the influence of demographic stochasticity on population growth rate. Dispersal also has serious consequences on population genetic structure (Cayuela et al. 2018a). An increase in dispersal (rates or distances) results in enhanced gene flow, which reduces the strength of genetic drift effects within subpopulations and homogenises allele frequencies across subpopulations (Broquet & Petit 2009). In addition, gene flow may strongly alter *N*_*e*_ estimates, the direction and the magnitude of the bias depending on the intensity, the continuity, and the randomness of gene flow (Wang & Whitlock 2003, Palstra & Ruzzante 2008). This led to the consideration of hierarchized *N*_*e*_ estimates, i.e. local *N*_*e*_ at the subpopulation level and meta-*N*_*e*_ at the whole population scale (Palstra & Ruzzante 2008). Therefore, considering the spatial structure of populations is a critical challenge for studies focusing on the contribution of demographic and genetic factors on extinction processes.

In this study, we examined demographic and genetic factors involved in the extinction process of a spatially structured population of a lekking bird, the western capercaillie (*Tetrao urogallus*). This species has experienced a severe decline in central and western Europe, mainly due habitat destruction and alteration (IUCN 2018). Populations of *T*. *urogallus* are usually composed of a set of discrete leks in which males actively compete for mates and between which dispersal occurs (Storch 1985, Wegge & Rolstad 1986, Cayuela et al. 2018b). Leks differ in terms of size (namely the number of competing males) and it has been reported in a related species with the same lekking behaviour that females preferentially reproduce in large leks (Alatalo et al. 1992). Using capture-recapture data and multievent capture-recapture models, we showed that the population experienced a dramatic decline over the study period (2010-2015). Then, we investigated the demographic mechanisms involved in this decrease. First, we examined if an Allee effect negatively affected adult survival and recruitment. Specifically, we expected (1) a lower recruitment and a reduced survival in small leks compared to large leks, and (2) we hypothesized that breeders could avoid a ‘mate finding’ Allee effects by adjusting their dispersal decisions according to lek size. We thus expected that adults (of both sexes) were more likely to emigrate from small leks and preferentially immigrated into large leks. Second, we examined how demographic–environmental stochasticity may affect survival and recruitment. We hypothesized that recruitment would show higher temporal variance than adult survival, which is a pattern usually observed in iteroparous species (Gaillard & Yoccoz 2003). In addition, we investigated whether survival and recruitment displayed a negative trend over time that could explain the population decline. Next, we examined potential genetic mechanisms (loss of genetic variation, decrease in effective population size and inbreeding depression) contributing to population decline. We hypothesized that the severe population loss was associated with a low allelic richness, a low expected heterozygosity, and high inbreeding coefficients (at both individual and population levels). We also expected that population decline would be associated with a small lek-specific *N*_*e*_, a phenomenon that should be amplified by the capercaillie mating system in which leks are mainly composed of related males and only a few dominant males reproduce (Höglund et al. 1999, Rintamäki et al. 2000, Kervinen et al. 2012, Cayuela et al. 2018b). In addition, as dispersal rates and gene flow are non-random and asymmetric in the population (Cayuela et al. 2018b, and **Supplementary material S3**), we expected that meta-*N*_*e*_ should be drastically lower than the sum of the local *N*_*e*_ (per lek) in the population. Last, we tracked evidence of inbreeding depression by examining if inbreeding negatively affected survival and recruitment.

## Material & Methods

### Study area and bird survey

The study was conducted in the Vosges Mountains, at the northern margin of *T*. *urogallus* distribution range in France (**Fig.1**). Large valleys structure the species distribution range into discrete forest patches. Euclidean distance between leks ranging from 2.6 to 42 km, with a median of 18 km. Non-invasive samples (95 % faeces and 5 % feathers) were collected from 2010 to 2015, which allowed us to identify 116 individuals (66 males and 50 females) distributed among eleven leks (**Supplementary material S1**, Table S1). Four historically large leks (NOI, LLC, VEN, and GdF) and seven small leks (TAN, CrH, FOS, TdR, BAG, GEH, and StA) were identified. During the first year of the study (2010), the maximum number of individuals captured in large leks ranged from 6 to 16 individuals, while in small leks it ranged from 1 to 6. A detailed description of the study area, the sampling method and the genotyping approach is described in a previous study (Cayuela et al. 2018b). The population was surveyed using the CR method over a 6-year period.

**Fig. 1.**
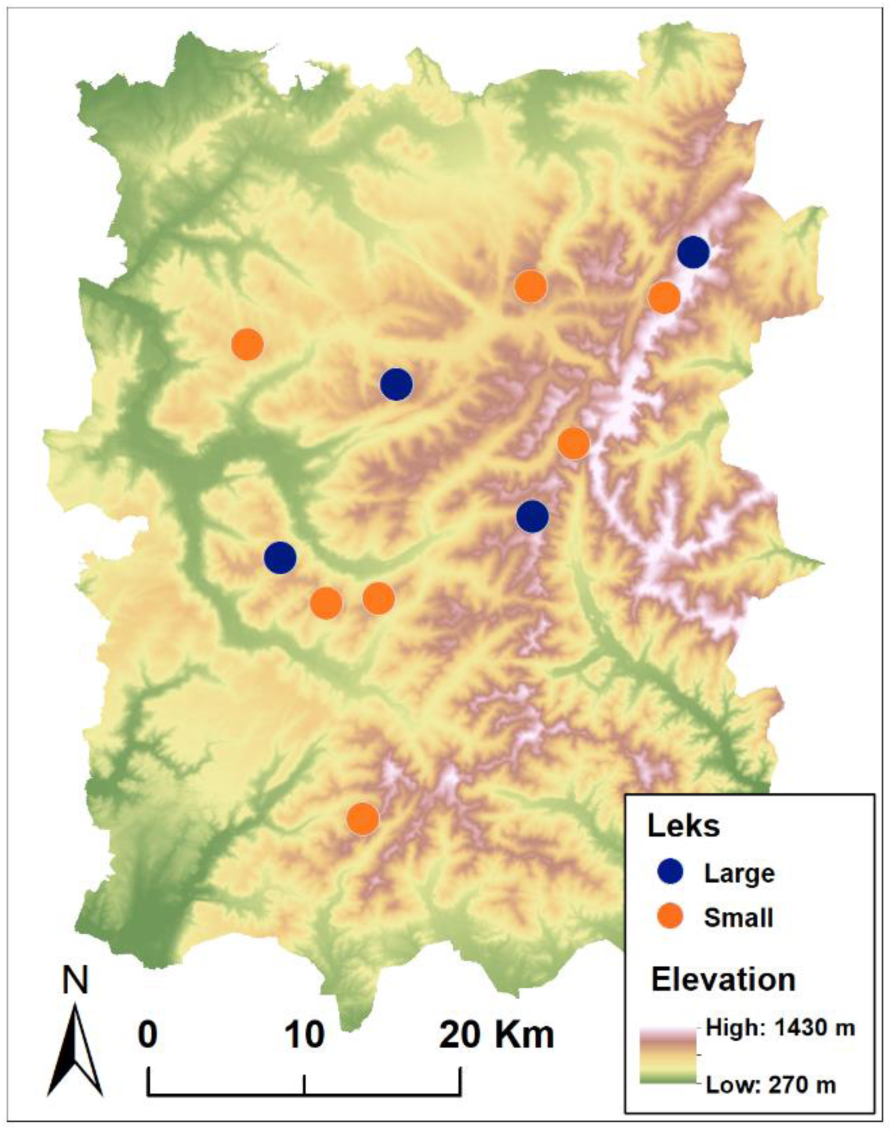
Map of the study area showing the eleven leks (four large and seven small) studied over the 6-years study period.

We analysed samples collected at lek sites during the breeding season (mid-March to mid-May) over a 6-year period using the CR method. We split samples collected during sampling visits into an early sampling session (S1), ranging from mid-March to mid-April, and a late sampling session (S2), ranging from mid-April to mid-May. Movements between leks during S1 were recorded as early dispersal events and, respectively, movements between leks occurring during S2 as late dispersal events.

### Modeling survival and dispersal according to lek size

Using multievent CR models, we examined how population size, survival and recruitment varied over time. We also investigated how lek size affects survival, dispersal and recruitment probability. In multi-event capture–recapture models, a distinction is made between events and states (Pradel 2005). An event is the field observation coded in the individual’s capture history, which is related to the latent state (e.g. alive or dead, dispersing or non-dispersing individuals) of the individual. These observations can result in a certain degree of uncertainty regarding the latent state. Multi-event models are designed to model this uncertainty in the observation process using hidden Markov chains (Pradel 2005).

We used a modified version (Tournier et al. 2017) of a multievent CR model proposed by Cayuela et al. (2017), which allows quantification of survival and dispersal between sites (in our case, leks) that vary spatially. The demographic parameters were modeled both intra-annually (i.e. between the two sessions S1 and S2 of the same year) and inter-annually. Our model had robust design structure (Pollock 1982) to quantify departure and arrival probabilities intra-annually and inter-annually. Mortality in adult capercaillie is highest in late winter and early spring, when capercaillie are among the rare available preys for avian and mammalian predators (Saniga 2011), and especially for males aggregated around lekking sites (Wegge et al. 1987). We therefore set the intra-annual survival probability (probability of survival from first to second sampling session) to 1, as is usually done in robust-design models (Kendall et al. 1995, 1997); this assumption was realistic in our study system as intra-annual survival probability was always higher than 0.9.

This model is based on states that include information about individual dispersal status between time *t*–1 and *t*, the capture status at *t*–1 and *t*, and the type of site (lek in our study system) occupied at *t*. This information is embedded in states as follows. Individuals that disperse between *t*–1 and t are coded ‘M’ for ‘moved’, while individuals that remain in the same lek are coded ‘S’ for ‘stayed’. Individual may occupy two different types of leks: small leks (‘P’) and large leks (‘L’). These codes are prefixed by the previous capture status and suffixed by the current capture status (+ for ‘captured’, o for ‘not captured’). This leads to the consideration of 13 states (described in **Table 2**). Six possible observations made in the field (i.e. events) are considered in the model. For individuals captured at *t* and *t*–1, a code of 1 or 4 was attributed if it did not change lek and occupied a small (code 1) or a large lek (code 4); 2 or 5 was attributed if it changed lek and occupied a small (code 2) or large lek (code 5). For individuals that were not captured at *t*–1 and were captured at *t* (so for which we had no information about whether or not they changed lek), we attributed a code of 3 if they were captured in a small lek or a code of 6 for a large lek. Individuals not captured at *t* were given a code of 0.

At its first capture, an individual could be in two states: oSP+ and oSL+. We then considered four state–state transition steps (described in the matrices presented in **Fig.2**): (1) survival, (2) departure, (3) arrival and (4) recapture. At the first modeling step, survival is updated and an individual may survive with a probability φ or die with a probability 1–ϕ. This leads to a matrix with 13 states of departure and seven states of arrival. Survival probability may depend on lek size (small ‘P’, or large ‘L’). Then, information about dispersal is updated and spread into two steps (2 and 3). At step 2, an individual may leave its lek with a probability ε or remain in the same lek with a probability 1–ε. Departure probability may also vary according to lek size. This results in a matrix with seven states of departure and 13 states of arrival. At step 3, an individual that has left its lek may arrive in a small lek with a probability α or in a large lek with a probability 1–α. This leads to a matrix with 13 states of departure and arrival. In step 4, information of individual capture is updated. Individuals may be captured with a probability *p* or not with a probability 1–*p*, resulting in a matrix of 13 states of departure and arrival. The last component of the model links events to states; each state corresponds to only one possible event (**Fig.2**).

**Fig. 2.**
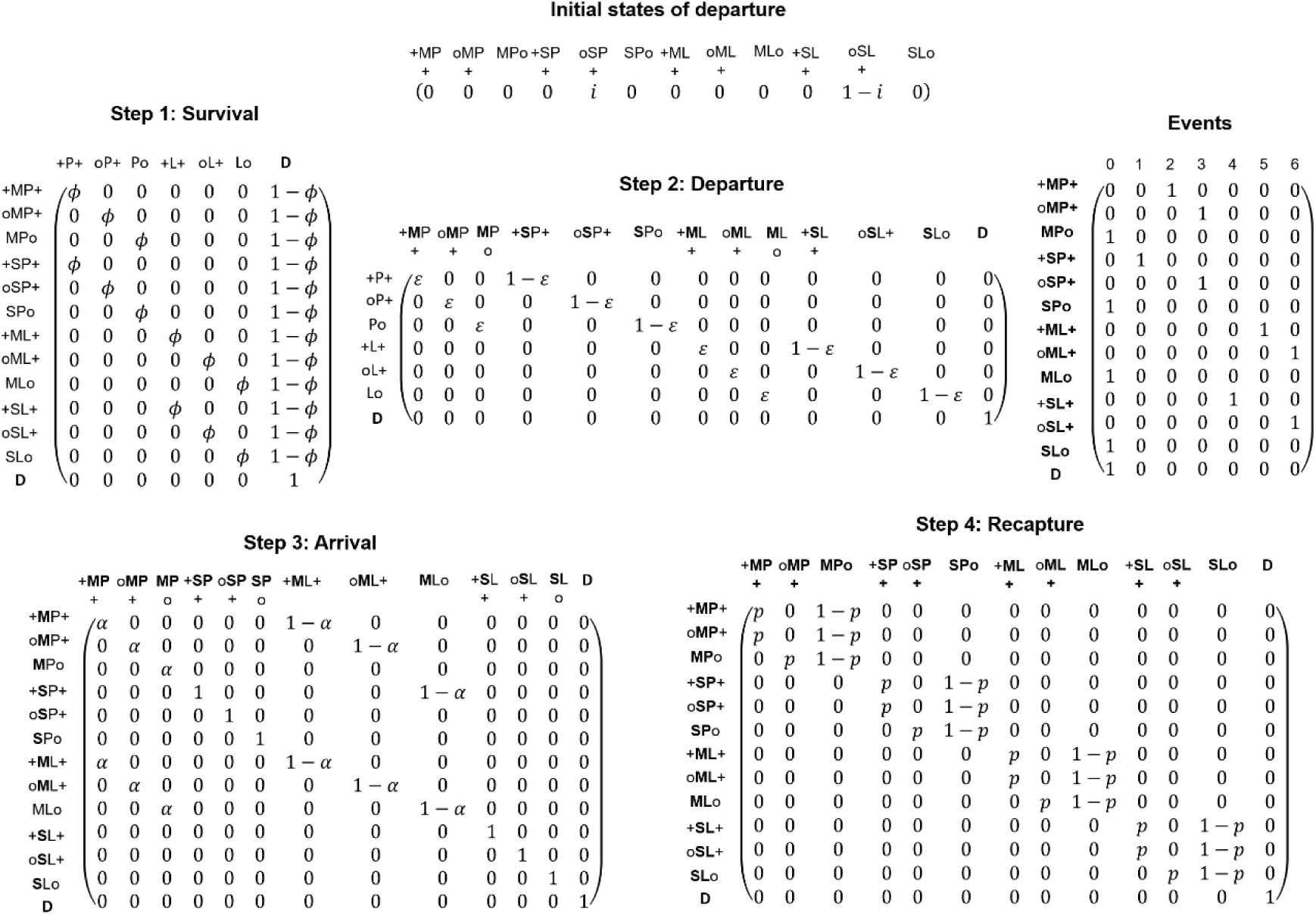
Matrices of the multievent model designed to estimate survival, dispersal (departure and arrival), and recapture probabilities in small and large leks. The model includes initial states of departure and four successive transition steps: (1) survival, (2) departure, (3) arrival, and (4) recapture. The matrix of events (described in Table 2) link field observations with model underlying states (described in Table 1). At each step, the information embedded in the composite state code was updated; the updated information appears in bold in the transition matrices.

**Table 1.** Description of the state of the multievent capture-recapture model.

| State | State description |
| --- | --- |
| +SP+ | Captured at $t-1$ and $t$ in the same small lek |
| +MP+ | Captured at $t$ in a small lek that is different from the lek where captured at $t-1$ |
| +SL+ | Captured at $t-1$ and $t$ in the same large lek |
| +ML+ | Captured at $t$ in a large lek that is different from the lek where captured at $t-1$ |
| oSP+ | Captured at $t$ in the same small lek as it occupied at $t-1$ when not captured |
| oMP+ | Captured at $t$ in a small lek that is different from the lek occupied at $t-1$ when not captured |
| oSL+ | Captured at $t$ in the same large lek as it occupied at $t-1$ when not captured |
| oML+ | Captured at $t$ in a large lek that is different from the lek occupied at $t-1$ when not captured |
| SPo | Not captured at $t$ and in the same small lek as at $t-1$ |
| MPo | Not captured at $t$ and in a small lek that is different from the lek occupied at $t-1$ |
| SLo | Not captured at $t$ and in the same large lek as at $t-1$ |
| MLo | Not captured at $t$ and in a large lek that is different from the lek occupied at $t-1$ |
| D | Dead |

This parameterization was entered in the program E-SURGE (Choquet et al. 2009). Competing models were ranked through a model-selection procedure using Akaike information criteria adjusted for a small sample size (AICc) and AICc relative weights (*w*). When the AICc weight of the best supported model was less than 0.9, we performed model averaging. The parameters were model-averaged using the complete set of models. Our hypotheses about recapture and state–state transition probabilities were examined using the general model [ϕ(SIZE + Y), τ(SIZE + SEX), α(SEX), *p*(SIZE + SEX + Y)], which included three effects: (1) lek size (SIZE) coded as states in the model; (2) year-specific variation (Y); (3) group effect for sex-specific variation (SEX). We examined whether recapture probability *p* differed according to lek size (SIZE), between years (Y) and between sexes (SEX). We tested whether survival depended on lek size (SIZE) and varied over years (Y). We did not examine the effect of sex (SEX) on survival as a previous study on this population revealed the absence of sex-specific variation in survival (Cayuela et al. 2018b). Concerning dispersal, we modeled the intra- and inter-annual departure rate and tested whether it varied according to lek size (SIZE) and between sexes (SEX). We did not consider time-specific variation in departure rate as dispersal rate displays little variation over time in this population (Cayuela et al. 2018b). Moreover, we examined whether arrival probability in small leks differed between sexes (SEX). All the combinations of these three effects were tested in the models, leading to the consideration of 64 competing models (**Supplementary material S2**). Moreover, we aimed to examine whether survival showed a negative trend over time. After determining the best-fitting model based on AICc and *w*, we examined the effect of time (included in the model as a continuous variable) on survival probability using ANODEV as recommended in Grosbois et al. (2008). Moreover, we were also interested in quantifying census population size (*N*_*b*_) per lek over the 6-year period. We estimated yearly population size by dividing the number of captured birds by the model-averaged recapture probability (i.e. Horvitz-Thompson estimator).

### Modeling recruitment according to lek size

We built a multi-event model to estimate recruitment following the structure of Pradel’s (1996) model, in which recruitment can be modelled by reversing capture histories and reading them backwards. In our model, the recruitment probability is estimated as the probability that an individual present at time *t* was not present at *t*–1, i.e. the proportion of “new” individuals in small and large leks at *t*, while taking into account dispersal between the two kinds of leks. The model has a relatively similar structure to that of the model designed to estimate survival. The two models have the same states and events, and the same modeling steps. The only feature that differs between the two models is the ‘survival’ matrix, which is replaced by a recruitment matrix in the recruitment model (**Fig.3**). At this modeling step, the information about recruitment is updated and an individual may be recruited with a probability δ or not with a probability 1–δ. This leads to a matrix with 13 states of departure and seven states of arrival; the recruitment probability may depend on lek size (small ‘P’, or large ‘L’)

**Fig. 3.**
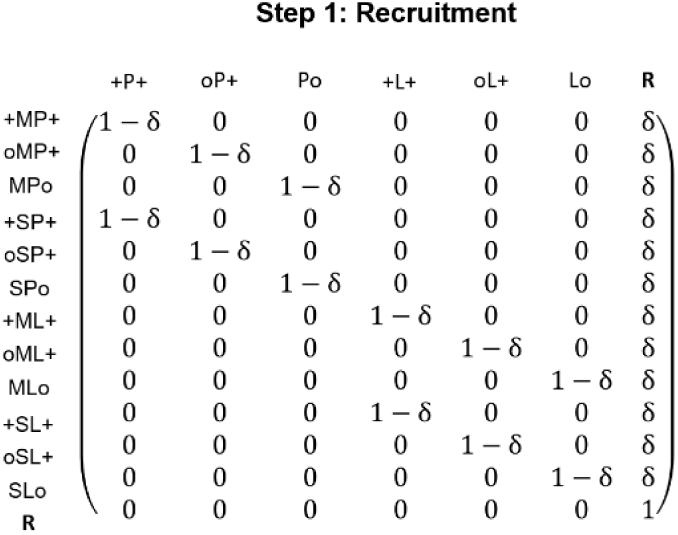
Recruitment matrix to estimate recruitment rate in small and large leks. The model has a similar structure to that of survival model (Fig.2). The survival matrix is replaced by a recruitment matrix with 13 states of departure and seven states of arrival.

This parameterization was implemented in the program E-SURGE. Competing models were ranked using AICc and AICc relative weights (w). We performed model averaging when the AICc weight of the best-supported model was less than 0.9. We examined our hypotheses about recapture and state–state transition probabilities from the general model [δ(SIZE + Y), τ(SIZE + SEX), α(SEX), *p*(SIZE + SEX + Y)], which includes the same effects that were considered in the survival model. As dispersal is not a biologically meaningful parameter in this models due to capture histories reversion, we kept the combination of effects, τ(SIZE + SEX) and α(SEX), in all the models. For recruitment and recapture, we tested all the possible combinations of effects, which led to the consideration of 16 competing models (**Supplementary material S2**).

### Quantifying allelic richness, inbreeding, gene flow and effective population size

For the genetic analyses, we used the dataset presented in Cayuela et al. (2018b). Ninety-three individuals were genotyped used 11 microsatellite markers (ADL184, ADL230, BG15, BG16, BG18, LEI098, TuT1, TuT2, TuT3, TuT4, ADL142). Evidence for scoring errors, large allele dropout, presence of null alleles, departure from Hardy–Weinberg equilibrium and linkage disequilibrium between all pairs of loci were previously assessed in Cayuela et al. (2018b).

Allelic richness (*A*, the uncorrected allelic richness; *A*_*r*_ the allelic richness corrected for rarefaction) and expected heterozygosity (*H*_*E*_) were calculated using FSTAT (Goudet 1995) for the four large leks and at the whole population level. We quantified *F*_*ST*_ and asymmetrical migration rates *m* between the four large leks using GENEPOP v4.1.4 (Rousset 2008) and BayesAss v1.3 (Wilson & Rannala 2003) respectively. BayesAss v1.3 is a bayesian method based on the temporary disequilibrium of genotypes (i.e., a process produced by immigration and attenuated by random mating), allowing the identification of recent immigrants and their offspring. This method does not assume migration–drift equilibrium, which is an assumption that is frequently violated in natural populations. We computed a total of 2,000,000 MCMC iterations after an initial burn-in of 1,000,000, as suggested by Wilson and Rannala (2003). We estimated effective population size (*N*_*e*_) using the program COLONY v2.0.6.4 (Jones & Wang 2010, Ackerman et al. 2017). We used default software options but set the mating system to polygamy in males and females and ‘Optimal Prior for Ne’ for sibship prior. Effective population sizes *N*_*e*_ were calculated at three hierarchical levels: (i) the lek level, (ii) at the level of four large leks, and (iii) at whole population level (i.e. meta*N*_*e*_). The ratio *N*_*e*_ /*N* were calculated by considering *N* as the population size at the beginning of the study (in 2010), i.e. prior to the rapid population loss recorded during the 6-year study period.

### Examining inbreeding influence on survival and recruitment

We estimated individual inbreeding coefficient using the *r* package Genhet (Coulon 2010). We considered three inbreeding metrics: (i) the proportion of heterozygous loci (PHt) in an individual, (ii) the internal relatedness (IR), and (iii) the homozygosity by locus (HL).

We examined the effects of the three inbreeding metrics on survival using Cormack-Jolly-Seber capture-recapture models. As we showed that recapture probability differed between sexes (see the result section), the sex was kept in all the models. We considered four alternative models in which survival varied according to PHt, IR or HL, or was kept constant (using the symbol ‘.’) (**Supplementary material S2**). The models were built in the program E-SURGE and were ranked using AICc and AICc relative weights (w). We used Pradel’s model (described above) to examine the effect of inbreeding on recruitment. The effect of the three inbreeding metrics was examined in a similar way as for survival.

## Results

### Population size, survival, recruitment and dispersal

Our analyses revealed that whole population size in the eleven leks dramatically dropped between 2010 and 2015 (**Fig.4**), from 136 to 90 individuals. It corresponds to a decrease of 33% of the population size in six years. This decline was much stronger in the large four leks (40%) than in small leks (20%) (**Fig.4**). The large leks GdF, VEN, LLC, and NOI lost 31%, 45%, 44%, and 50% of their size respectively.

The best-supported survival model was [ϕ(SIZE), τ(SIZE), α(.), *p*(SEX)] (see the complete model selection procedure in Supplementary material S2, Table S1). However, as its AICc weight was 0.27, we performed model-averaging. Recapture probability differed between sexes, males (0.58±0.03) having a higher recapture probability than females (0.37±0.04). Survival showed little variation over time and was marginally higher in small leks (from 0.70 in 2013 to 0.72 in 2015) than in large leks (from 0.65 in 2013 to 0.68 in 2015) (**Fig.5A**). Our study revealed that dispersal was strongly affected by lek size. Departure probability was drastically higher in small leks than in large leks (**Fig.5C**). In both males and females, the probability of leaving a small leks was 0.25±0.07 inter-annually and 0.25±0.07 intra-annually, while it was 0.14±0.04 inter-annually and 0.15±0.03 intra-annually in large leks. Simultaneously, individuals had a higher probability to arrive in a large lek than in a small one (**Fig.5D**). Intra-annually, the probability of arriving in a large lek was 0.65±0.10 while it was 0.35±0.10 in a small lek. This pattern was less marked inter-annually: the probability of arriving in a large lek was 0.55±0.12 while it was 0.44±0.12 in a small lek.

**Fig. 4.**
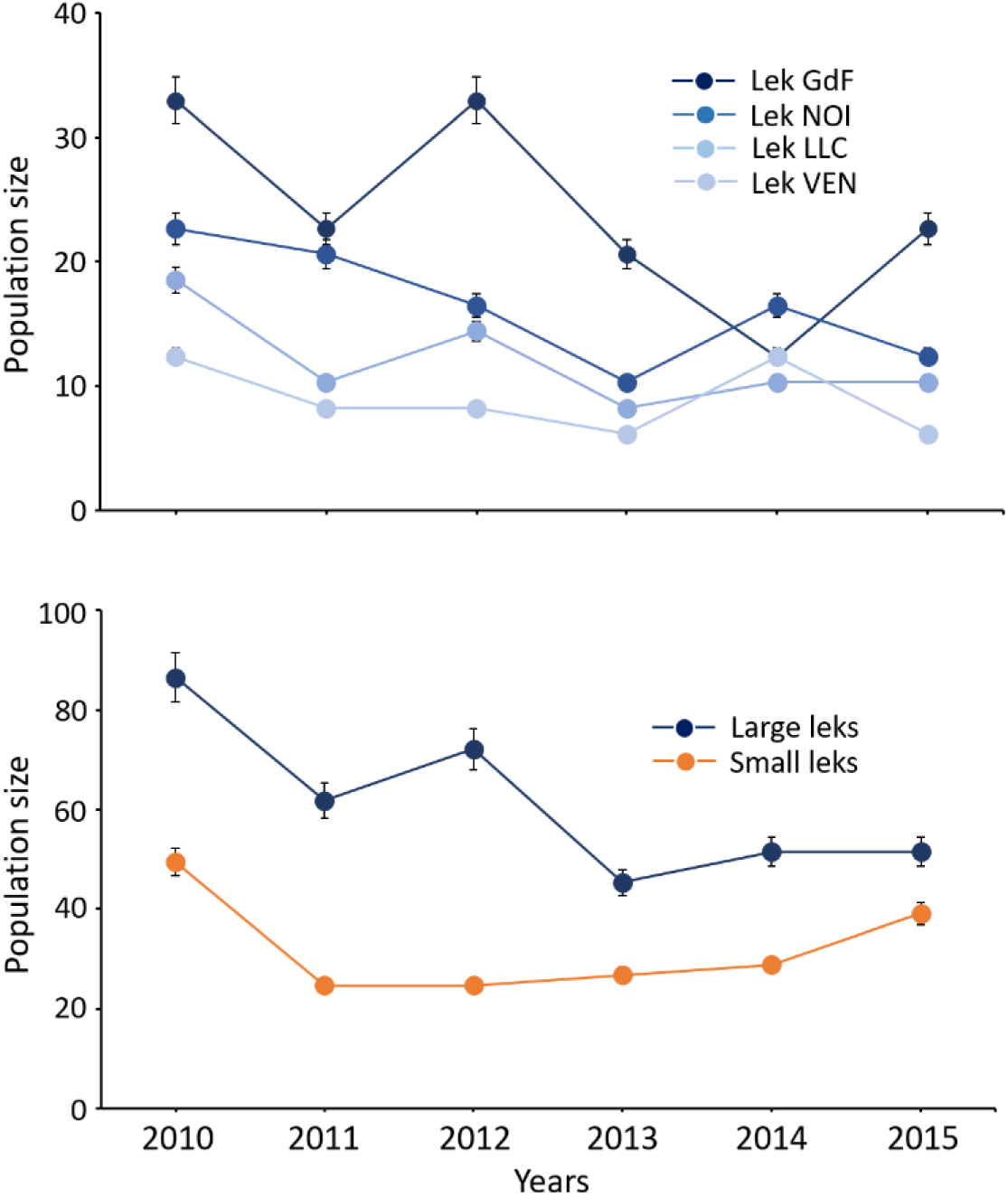
Decline of the population of western capercaillie (*Tetrao urogallus*) over the 6-years study period (2010-2015). (A) annual size of the four large leks. (B) annual size of large and small leks.

**Fig. 5.**
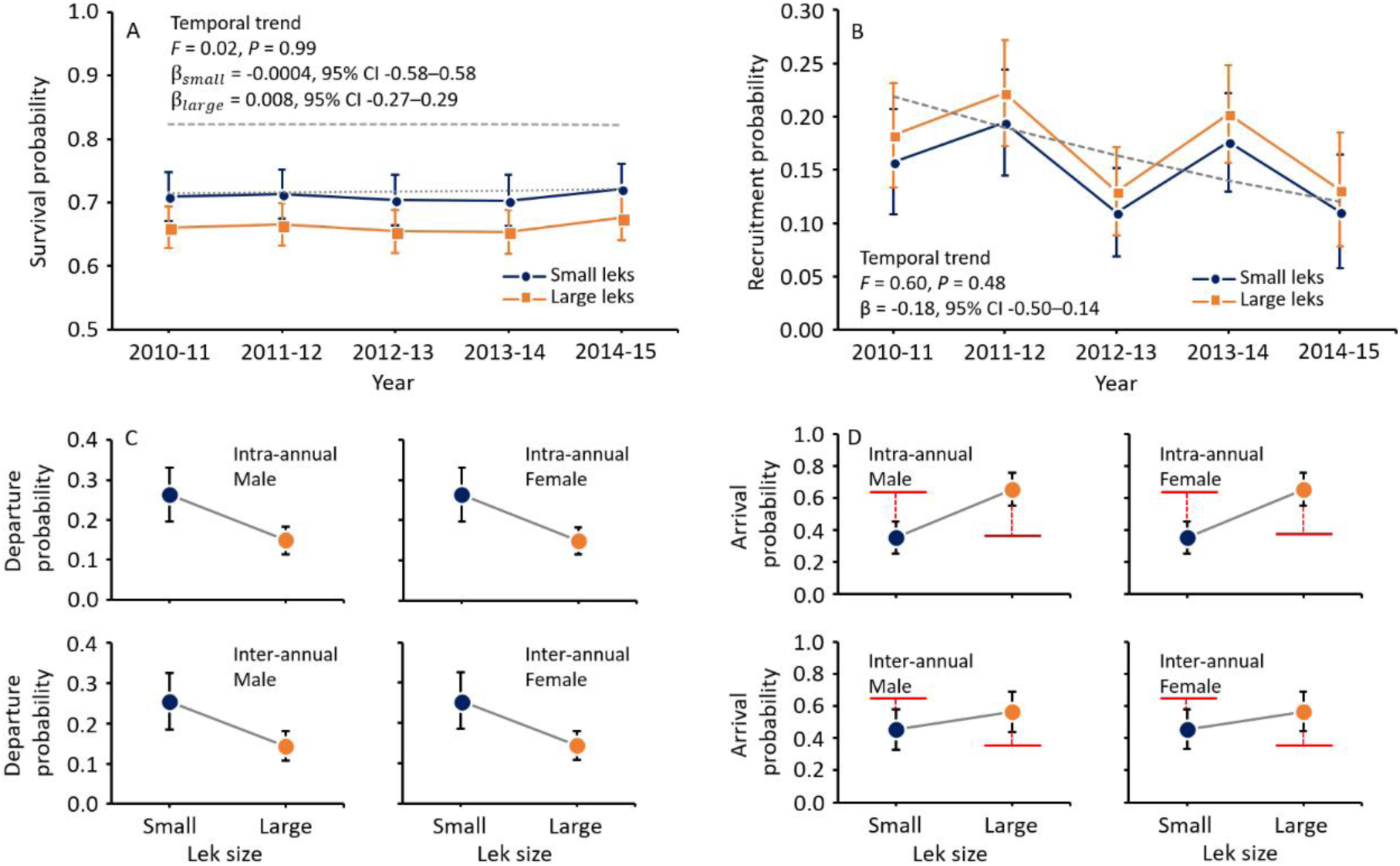
Survival, recruitment and dispersal in a declining population of western capercaillie (*Tetrao urogallus*) over a 6-year study period (2010-2015). Model-averaged estimates and their standard errors are extracted from multievent capture-recapture models. Survival (A) marginally differs between large and small leks and shows little temporal variation. Survival does not show any significant temporal trend (small leks = grey dashed line, large leks = grey dotted line). Recruitment (B) does not vary between large and small leks. It shows substantial variation over time but no significant temporal trend (grey dashed line). Dispersal (C-D) strongly depends on lek size. Departure probability (C) is lower in large leks than in small leks. By contrast, arrival probability is higher in large leks than in small leks. The red bar shows the probability of arriving in a lek (small or large) under random expectation.

The best-supported recruitment model was [δ(.), τ(SIZE + SEX), α(SEX), *p*(SEX)] (see the procedure of model selection in Supplementary material S2, Table S2). However, as its AICc weight was 0.18, model-averaging was performed. Recruitment probability did not differ between small and large leks (**Fig.5B**). Recruitment probability varies over time (**Fig.5B**), ranging from 0.11±0.04 in 2013 and 2015 to 0.19±0.05 in 2012.

### Allelic richness, inbreeding, gene flow and effective population size

The allelic richness corrected for rarefaction ranged from 2.23 in GdF and LCC to 2.70 in NOI (**Fig.6**). Those values were relatively close to allelic richness calculated for the four main leks (2.61) and the whole population (2.64). The expected heterozygosity (*H*_*e*_) ranged from 0.39 in GdF to 0.46 in NOI. It was 0.44 at the large lek and the whole population levels. Inbreeding coefficients (*F*_*IS*_) ranged from −0.01 in VEN to 0.11 in NOI and GdF (**Fig.6**). Concerning the individual metrics of inbreeding, the proportion of heterozygous loci (PHt) was 0.37±0.17 (min = 0, max = 0.80), the internal relatedness (IR) was 0.17±0.34 (min = −0.53, max = 1), and the homozygosity by locus (HL) was 0.56±0.20 (min = 0.14, max = 1). In the four large leks, the mean *F*_*IS*_ was 0.17 while it 0.14 in the whole population. Moreover, *F*_*ST*_ values between the four large leks ranged from 0.08 to 0.21 (**Fig.6**). Asymmetrical migration rates *m* between the four large leks ranged from less than 0.01 to 0.12 (**Fig.6**).

**Fig. 6.**
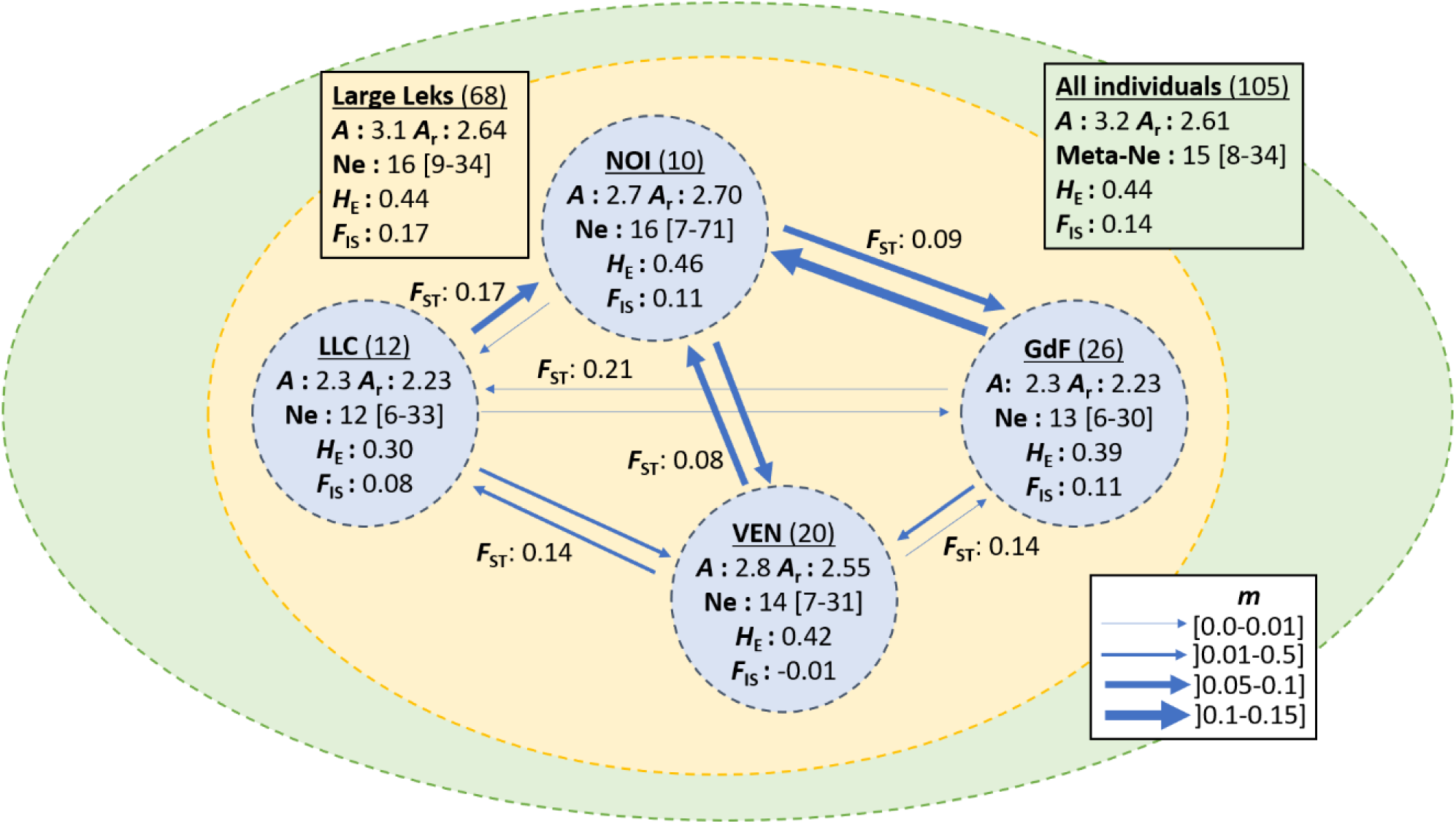
Effective population size (*N*_*e*_ for the four large leks, and meta-*N*_*e*_ for the whole population), asymmetrical migration rate (*m*), allelic richness, and inbreeding coefficient (*F*_*IS*_) in a declining population of western capercaillie (*Tetrao urogallus*). For the four large leks, we show estimates of *N*_*e*_, *m, F*_*IS*_, allelic richness (*A*, uncorrected richness; *A*_*r*_ richness corrected for rarefaction), and expected heterozygosity (*H*_*E*_). We also give *F*_*ST*_ values for each pair of large leks.

The estimates of *N*_*e*_ were relatively similar in the four large leks (**Fig.6**), ranging from 12 (95% CI 3–33) in LLC to 16 (95% CI 7–71) in NOI. Furthermore, the mean *N*_*e*_ in the four leks was 16 (95% CI 9–34). The meta-*N*_*e*_ estimated for the whole population was 15 (95% CI 8–34). The ratios *N*_*e*_ /*N* was always lower than 1 in the large leks (VEN = 0.93, LCC = 0.67, GdF = 0.52; not estimated for NOI) and at whole population level (0.15).

### Examining inbreeding influence on survival and recruitment

The model with constant survival [ϕ(.), *p*(SEX)] was better supported than the models including one of three inbreeding metrics (Supplementary material S2, Table S3). Moreover, the slope coefficient of inbreeding metrics was close to 0 (PHt: −0.06±0.16; IR: 0.06±0.16; HL: 0.01±0.16), indicating a very small size effect. Overall, this indicates that survival was weakly affected by individual inbreeding regardless of the inbreeding metric considered.

Similarly, the model with constant recruitment [δ (.), *p*(SEX)] was better supported than the models incorporating the three inbreeding metrics (Supplementary material S2, Table S4). Furthermore, the slope coefficient of each metric was different of 0, (PHt: 0.21; IR: −0.39; HL: - 0.25) but the confidence intervals always included 0, thus suggesting a low accuracy of model estimates (PHt: −0.40–0.81; IR: −1.09–0.30; HL: −0.91–0.41). Therefore, inbreeding seemed to have a little influence on recruitment regardless of the metrics used to quantify inbreeding.

## Discussion

Our study showed that the population of *T*. *urogallus* experienced a severe decline between 2010 and 2015. We did not detect any Allee effect on survival and recruitment; by contrast, a behavioral response to a mate finding Allee effect was found, individuals of both sexes dispersing to avoid small leks. Survival was relatively constant over time, while recruitment was more variable. In parallel to this demographic decline, the population displayed a low genetic diversity and a high inbreeding (compared to that reported in this species, Segelbacher et al. 2002, 2003; Rodríguez-Muñoz et al. 2007, Klinga et al. 2015). The effective population size at both lek and population levels was low. Furthermore, we did not find evidence of inbreeding depression as inbreeding affected neither survival nor recruitment probability.

### Demographic processes

Our study revealed a dramatic population decline (−52%) over the 6-year period. The decline experienced by the population is likely caused by a loss of recruitment rather than decrease of adult survival; recruitment does not compensate the ‘normal’ adult mortality which leads to population decline. Indeed, the adult survival in the studied population was similar to that reported in other population of *T*. *urogallus* (Storch 2007 and references therein). The low temporal variation of survival probability indicates limited demographic and environmental stochasticity for this demographic parameter. This pattern is commonly found in relatively long-lived species in which selection favors phenotypic traits that reduce the temporal variance of adult survival (Gaillard & Yoccoz 2003). Moreover, our analyses did not reveal any temporal trend for survival that could explain the population decline. By contrast, recruitment rate was more variable between years, with substantial drops in 2013 and 2015. The trend analysis did not reveal any statistically significant gradual changes of recruitment over time. However, the negative coefficient slope of time effect in the model suggests that we failed to detect a negative trend, probably due to a lack of statistical power; the small population size and the shortness of study period likely decreases model estimate accuracy.

The cause of population decline remains unclear so far. The population loss was stronger in the large leks (−40%) in the small ones (−20%). This pattern indicates the absence of an Allee effect on population growth (called ‘demographic Allee effect’, Stephens et al. 1999); population growth is lower in large leks than in small leks. In addition, our results also indicate that survival and recruitment are not affected by lek size, which also suggest the absence of Allee effect on fitness components. By contrast, dispersal probabilities (both departure and arrival) were non-random and indicated a behavioral response to ‘mate finding Allee effect’ (i.e. difficulty to locate mate due to low conspecific density). Both males and females had a lower probability of emigrating from large leks; in parallel, dispersers of both sexes more often immigrated into large leks. In tetraonids, females preferentially reproduce in large leks, which results in a higher reproductive success for the males attending these leks (Alatalo et al. 1992). Aggregation of individuals likely reduce the demographic Allee effect and limit the effects of demographic stochasticity in the largest leks. By contrast, it reduces the chance of new lek formation and enhances the risk of small lek disappearance (low ‘rescue effect’, Hanski et al. 1997).

The high variation of recruitment over time (and possibly over space) suggests that intrinsic and/or extrinsic factors affecting the recruitment process were the main drivers of population decline. Our study shows that recruitment variation involved was likely independent from lek size. However, environmental factors such as habitat loss and alteration could have a detrimental effect on female fecundity and chick survival, and therefore on adult recruitment few years later. Moss et al. (2008) showed that gradual shift from early to mid-April of spring warming reduced synchrony between the start of vegetation growth and peak of juvenile energetic demands, and potentially decreased female reproductive success. Rapid change in vegetation phenology in the Vosges Mountains also correlated with delayed peak of activity at capercaillie lek and suggests a negative impact of climatic factors on recruitment rate in this population (Ménoni et al. 2012). Genetic factors could also cause recruitment variation.

### Genetic processes

Our study highlighted low genetic diversity and high inbreeding in this declining population. The mean allelic richness (2.64 at the whole population level), and the expected heterozygosity (0.44) are low compared to those reported in *T*. *urogallus* populations from central and northern Europe and relatively similar to those measured in small, fragmented populations (Segelbacher et al. 2002, 2003; Rodríguez-Muñoz et al. 2007). The inbreeding coefficient *F*_*IS*_ (0.14) was among the highest reported in the range of *T*. *urogallus* (Segelbacher et al. 2002, 2003; Rodríguez-Muñoz et al. 2007, Klinga et al. 2015), close to the *F*_*IS*_ of 0.15 reported in a population from western Carpathians (Klinga et al. 2015). Fragmentation of the distribution range of the species during the 20^th^ century (Storch 2007) led to the loss of genetic connectivity between the Alpine core population and peripheral, small populations, as described by Segelbacher et al (2003). At that time, *i*.*e*. approximately 5–6 generations ago, the authors already reported high levels of genetic differentiation between the population in the Vosges Mountains and the Alpine population (*F*_*ST*_ > 0.10). It is thus very likely that the level of genetic differentiation between these two populations further increased, exacerbated by the decline of the population and the low lek-specific *N*_*e*_.

At the lek level, the ratio *N*_*e*_ /*N* < 1 results from the mating system of *T*. *urogallus*, in which few dominant males monopolize most of the mates, leading to a skewed reproductive success. This pattern is congruent with results from a previous study focusing on another grouse species with a lekking behaviour (Stiver et al. 2008). At the whole population level, the meta-*N*_*e*_ is far less than the sum of the lek-specific *N*_*e*_. This likely results from a non-random dispersal (depending on lek size) that should result in asymmetric gene flow between small and large leks. Migration rates *m* between large leks are also asymmetric, which suggests that other social factors than lek size affects dispersal. A previous study on this population thus found that females disperse in response to inbreeding risks and males preferentially join leks composed of relatives (Cayuela et al. 2018b). Furthermore, landscape resistance (especially landform) also results in asymmetric gene flow in the population (**Supplementary material S3**). These results emphasize the importance of considering dispersal patterns and population genetic structuring for the estimation and the interpretation of *N*_*e*_ (Wang & Whitlock 2003, Palstra & Ruzzante 2008).

Although we highlighted a high inbreeding level at both lek and population scales, we did not find evidence of inbreeding depression at the adult stage in our study system. Both survival and recruitment of adults were not affected by individual inbreeding. Given the small size of the population, one cannot rule out the possibility that the absence of significant effect was due to a low statistical power. That being said, for survival, the slope of coefficients associated with the inbreeding metrics were very close to 0, which suggests a small size effect rather than a low accuracy of the estimates. For recruitment, the accuracy of model estimates was low, likely to due to our small sample size. Inbreeding has been reported to negatively affect egg hatchability rate in the Greater prairie chicken (Westemeier et al. 1998), susceptibility to parasite and juvenile mortality in capercaillie (Isomursu et al. 2012) and male lifetime copulation success Black grouse (Höglund et al. 2002), no detrimental effect of inbreeding on survival at adult stages has already been reported in those birds. Future studies should investigate inbreeding effects on survival and females’ fecundity in large populations to increase the statistical power of the analyses. To conclude, we do not have any evidence, in our study system or in others, that the high inbreeding level has a detrimental impact of survival-related or recruitment-related performances in *T*. *urogallus*.

### Relative effects of demographic and genetic processes

The rapid decline of the population (50% of loss in six years, i.e. in less than two generations) suggests a higher contribution of the demographic factors than the genetic ones in the decline of the population. This interpretation is congruent with the statements of Lande (1988; and others later, Elgar & Clode 2001, Wootton & Pfister 2013) who postulated that demographic factors usually act faster than genetic ones in biological extinction processes. Although we did not highlight inbreeding effect on adult survival and recruitment, it remains nevertheless possible that inbreeding speeds up the population decrease, by affecting female fecundity, egg hatchability, and chick survival – three parameters not considered in this study and which also contribute to adult recruitment. However, in lek-breeding species, mechanisms related to sexual selection (i.e. disassortative mating or heterozygosity-based mate choice; Tregenza & Wedell 2000, Ryder et al. 2009) and dispersal (i.e. context-dependent dispersal based on inbreeding avoidance, Lebigre et al. 2010, Cayuela et al. 2018b) may limit the risk of inbreeding. Future studies should be undertaken to better understand if and how these behavioural mechanisms allow mitigating inbreeding depression and the contribution of genetic factors in biological extinctions.

## Supporting information

Sup Mat1

Sup Mat2

Sup Mat3

## Acknowledgment

The genetic monitoring of capercaillie in the Vosges Mountains was funded by the Life+ Project “Des Forêts pour le Grand tetras”, by the Natura2000 network and by the regional programme of the Capercaillie National Action Plan initiated by the French Ministry of Environment. The project largely relied on the work of volunteers who collected samples during the six years of the study: Antoine Andre, Didier Arseguel, Samuel Audinot, Alix Badre, Etienne Barbier, Dominique Becker, Bernard Binetruy, Frédéric Bocquenet, Noémie Castaing, Sebastien Coulette, Stéphane Damervalle, Luc Dauphin, Richard Delaunay, Lucile Demaret, Michel Despoulin, Laurent Domergue, Vincent Drillon, Christian Dronneau, Fabien Dupont, Arnaud Foltzer, Patrick Foltzer, Marc Gehin, Cyril Gerard, Maxime Girardin, Remi Grandemange, Jean-Claude Gregy, Joaquim Hatton, Thibaut Hingray, Thierry Hue, Arnaud Hurstel, Jean-Nöel Journot, Fabien Kilque, Lydie Lallement, Christian Lamboley, Manuel Lembke, Jean-Michel Letz, Vincent Lis, Olivier Marchand, Paul Massard, Yvan Mougel, Michel Munier, Louis-Michel Nageleisen, Yvan Nicolas, Christian Oberle, Pascal Perrotey-Doridant, Christian Philipps, François Rey-Demaneuf, Dorian Toussaint, Jean-Marie Triboulot, Bruno Vaxelaire, Laurent Verard, Jean-Lou Zimmermann, and Alice Zimmermann.

## References

Ackerman, M. W., Hand, B. K., Waples, R. K., Luikart, G., Waples, R. S., Steele, C. A., Garner, B. A., McCane, J., & Campbell, M. R. (2017). Effective number of breeders from sibship reconstruction: Empirical evaluations using hatchery steelhead. Evolutionary applications, 10(2), 146–160.

Alatalo, R. V., Höglund, J., Lundberg, A., & Sutherland, W. J. (1992). Evolution of black grouse leks: female preferences benefit males in larger leks. Behavioral Ecology, 3(1), 53–59.

Ballou, J., & Ralls, K. (1982). Inbreeding and juvenile mortality in small populations of ungulates: a detailed analysis. Biological Conservation, 24(4), 239–272.

Benton, T. G., & Bowler, D. E. (2012). Linking dispersal to spatial dynamics. Dispersal ecology and evolution, 251–265.

Briskie, J. V., & Mackintosh, M. (2004). Hatching failure increases with severity of population bottlenecks in birds. Proceedings of the National Academy of Sciences, 101(2), 558–561.

Broquet, T., & Petit, E. J. (2009). Molecular estimation of dispersal for ecology and population genetics. Annual Review of Ecology, Evolution, and Systematics, 40, 193–216.

Cayuela, H., Pradel, R., Joly, P., & Besnard, A. (2017). Analysing movement behaviour and dynamic space-use strategies among habitats using multi-event capture-recapture modelling. Methods in Ecology and Evolution, 8(9), 1124–1132.

Cayuela, H., Rougemont, Q., Prunier, J. G., Moore, J. S., Clobert, J., Besnard, A., & Bernatchez, L. (2018a). Demographic and genetic approaches to study dispersal in wild animal populations: a methodological review. Molecular ecology. In press.

Cayuela*, H., Boualit*, L., Laporte, M., Prunier, J. G., Foletti, F., Clobert, J., Jacob, G. (2018b) Kin-dependent dispersal influences relatedness and genetic structuring in a lek system. Oecologia, accepted.

Choquet, R., Rouan, L., & Pradel, R. (2009). Program E-SURGE: a software application for fitting multievent models. In: Modeling demographic processes in marked populations. Thomson, D. L., Cooch, E. G., Conroy, M. J. (eds). pp. 845–865. Springer, US.

Clobert, J., Le Galliard, J. F., Cote, J., Meylan, S., & Massot, M. (2009). Informed dispersal, heterogeneity in animal dispersal syndromes and the dynamics of spatially structured populations. Ecology letters, 12(3), 197–209.

Coulon, A. (2010). GENHET: an easy-to-use R function to estimate individual heterozygosity. Molecular Ecology Resources, 10(1), 167–169.

Courchamp, F., Clutton-Brock, T., & Grenfell, B. (1999). Inverse density dependence and the Allee effect. Trends in ecology & evolution, 14(10), 405–410.

Crnokrak, P., & Roff, D. A. (1999). Inbreeding depression in the wild. Heredity, 83(3), 260.

Elgar, M. A., & Clode, D. (2001). Inbreeding and extinction in island populations: a cautionary note. Conservation Biology, 15(1), 284–286.

Engen, S., Lande, R., & SÆther, B. E. (2003). Demographic stochasticity and Allee effects in populations with two sexes. Ecology, 84(9), 2378–2386.

Engen, S., Lande, R., & Saether, B. E. (2005). Effective size of a fluctuating age-structured population. Genetics.

Frankham, R., Ballou, J.D., Briscoe, D.A., 2002. Introduction to Conservation Genetics. Cambridge University Press, Cambridge.

Frankham, R. (2005). Genetics and extinction. Biological conservation, 126(2), 131–140.

Gaillard, J. M., & Yoccoz, N. G. (2003). Temporal variation in survival of mammals: a case of environmental canalization?. Ecology, 84(12), 3294–3306.

Gascoigne, J., Berec, L., Gregory, S., & Courchamp, F. (2009). Dangerously few liaisons: a review of mate-finding Allee effects. Population Ecology, 51(3), 355–372.

Gibbs, J. P. (2001). Demography versus habitat fragmentation as determinants of genetic variation in wild populations. Biological Conservation, 100(1), 15–20.

Gilpin, M. (Ed.). (2012). Metapopulation dynamics: empirical and theoretical investigations. Academic Press.

Goudet, J. (1995). FSTAT (version 1.2): a computer program to calculate F-statistics. Journal of heredity, 86(6), 485–486.

Grosbois, V., Gimenez, O., Gaillard, J. M., Pradel, R., Barbraud, C., Clobert, J., Grosbois, V., Gimenez, O., Gaillard, J. M., Pradel, R., Barbraud, C., Clobert, J., Møller, A. P. & Weimerskirch, H. (2008). Assessing the impact of climate variation on survival in vertebrate populations. Biological Reviews, 83(3), 357–399.

Hanski, I., Gilpin, M. E., & McCauley, D. E. (1997). Metapopulation biology (Vol. 454). San Diego: Academic press.

Hill, M. F., Hastings, A., & Botsford, L. W. (2002). The effects of small dispersal rates on extinction times in structured metapopulation models. The American Naturalist, 160(3), 389–402.

Höglund, J., Alatalo, R. V., Lundberg, A., RintamÎki, P. T., & Lindell, J. (1999). Microsatellite markers reveal the potential for kin selection on black grouse leks. Proceedings of the Royal Society of London B: Biological Sciences, 266(1421), 813–816.

Höglund, J., Piertney, S. B., Alatalo, R. V., Lindell, J., Lundberg, A., & Rintamäki, P. T. (2002). Inbreeding depression and male fitness in black grouse. Proceedings of the Royal Society of London B: Biological Sciences, 269(1492), 711–715.

IUCN 2018. The IUCN Red List of Threatened Species. Version 2018-1.

Isomursu M, Rätti O, Liukkonen-Anttila T, Helle P. (2012). Susceptibility to intestinal parasites and juvenile survival are correlated with multilocus microsatellite heterozygosity in the Capercaillie (Tetrao urogallus). Ornis Fennica 89:109–119

Jaquiéry, J., Guillaume, F., & Perrin, N. (2009). Predicting the deleterious effects of mutation load in fragmented populations. Conservation Biology, 23(1), 207–218.

Jones, O. R., & Wang, J. (2010). COLONY: a program for parentage and sibship inference from multilocus genotype data. Molecular Ecology Resources, 10(3), 551–555.

Keller, L. F. (1998). Inbreeding and its fitness effects in an insular population of song sparrows (Melospiza melodia). Evolution, 52(1), 240–250.

Kendall, W. L., Pollock, K. H., & Brownie, C. (1995). A likelihood-based approach to capture-recapture estimation of demographic parameters under the robust design. Biometrics, 293–308.

Kendall, W. L., Nichols, J. D., & Hines, J. E. (1997). Estimating temporary emigration using capture–recapture data with Pollock’s robust design. Ecology, 78(2), 563–578.

Kervinen, M., Alatalo, R. V., Lebigre, C., Siitari, H., & Soulsbury, C. D. (2012). Determinants of yearling male lekking effort and mating success in black grouse (Tetrao tetrix). Behavioral Ecology, 23(6), 1209–1217.

Klinga, P., Mikoláš, M., Zhelev, P., Höglund, J., & Paule, L. (2015). Genetic differentiation of western capercaillie in the Carpathian Mountains: the importance of post glacial expansions and habitat connectivity. Biological Journal of the Linnean Society, 116(4), 873–889.

Lande, R. (1988). Genetics and demography in biological conservation. Science, 241(4872), 1455–1460.

Lande, R. (1993). Risks of population extinction from demographic and environmental stochasticity and random catastrophes. The American Naturalist, 142(6), 911–927.

Lande, R. (1998). Anthropogenic, ecological and genetic factors in extinction and conservation. Researches on population ecology, 40(3), 259–269.

Lande, R., & Orzack, S. H. (1988). Extinction dynamics of age-structured populations in a fluctuating environment. Proceedings of the National Academy of Sciences, 85(19), 7418–7421.

Landguth, E. L., Muhlfeld, C. C., Waples, R. S., Jones, L., Lowe, W. H., Whited, D., Lucotch, J., Neville, H., & Luikart, G. (2014). Combining demographic and genetic factors to assess population vulnerability in stream species. Ecological Applications, 24(6), 1505–1524.

Lebigre, C., Alatalo, R. V., & Siitari, H. (2010). Female-biased dispersal alone can reduce the occurrence of inbreeding in black grouse (Tetrao tetrix). Molecular Ecology, 19, 1929–1939.

Luque, G. M., Vayssade, C., Facon, B., Guillemaud, T., Courchamp, F., & Fauvergue, X. (2016). The genetic Allee effect: a unified framework for the genetics and demography of small populations. Ecosphere, 7(7).

Mathieu-Bégné, E., Loot, G., Chevalier, M., Paz-Vinas, I., Blanchet, S. (2018). Demographic and genetic collapses in spatially structured populations: insights from a long-term survey in wild fish metapopulations. Oikos.

Matthysen, E. (2012). Multicausality of dispersal: a review. Dispersal ecology and evolution, 27, 3–18.

Meagher, S., Penn, D. J., & Potts, W. K. (2000). Male–male competition magnifies inbreeding depression in wild house mice. Proceedings of the National Academy of Sciences, 97(7), 3324–3329.

Melbourne, B. A., & Hastings, A. (2008). Extinction risk depends strongly on factors contributing to stochasticity. Nature, 454(7200), 100.

Ménoni E, Montadert M, Leclercq B, Hurstel A, Dillet K. 2012. Change in mating and breeding time of the Capercaillie in France, in relation to the change of the phenology of spring vegetation, 12th International Grouse Symposium, Matsumoto, 21–24th July 2012.

Moss R, Oswald J, Baines D. 2008. Climate change and breeding success: decline of the capercaillie in Scotland. Journal of Animal Ecology, 70, 47–61.

Naeem, S., Duffy, J. E., & Zavaleta, E. (2012). The functions of biological diversity in an age of extinction. Science, 336(6087), 1401–1406.

Nunney, L., & Campbell, K. A. (1993). Assessing minimum viable population size: demography meets population genetics. Trends in Ecology & Evolution, 8(7), 234–239.

O’Grady, J. J., Brook, B. W., Reed, D. H., Ballou, J. D., Tonkyn, D. W., & Frankham, R. (2006). Realistic levels of inbreeding depression strongly affect extinction risk in wild populations. Biological Conservation, 133(1), 42–51.

Palstra, F. P., & Ruzzante, D. E. (2008). Genetic estimates of contemporary effective population size: what can they tell us about the importance of genetic stochasticity for wild population persistence?. Molecular Ecology, 17(15), 3428–3447.

Purvis, A., Gittleman, J. L., Cowlishaw, G., & Mace, G. M. (2000). Predicting extinction risk in declining species. Proceedings of the Royal Society of London B: Biological Sciences, 267(1456), 1947–1952.

Pimm, S. L., Jenkins, C. N., Abell, R., Brooks, T. M., Gittleman, J. L., Joppa, L. N., Raven, P. H., Roberts, C. M., & Sexton, J. O. (2014). The biodiversity of species and their rates of extinction, distribution, and protection. Science, 344(6187), 1246752.

Pollock, K. H. (1982). A capture-recapture design robust to unequal probability of capture. The Journal of Wildlife Management, 46(3), 752–757.

Pradel, R. (1996). Utilization of capture-mark-recapture for the study of recruitment and population growth rate. Biometrics, 703–709.

Pradel, R. (2005). Multievent: an extension of multistate capture–recapture models to uncertain states. Biometrics, 61(2), 442–447.

Reed, D. H., & Frankham, R. (2001). How closely correlated are molecular and quantitative measures of genetic variation? A meta-analysis. Evolution, 55(6), 1095–1103.

Reed, D. H. (2005). Relationship between population size and fitness. Conservation biology, 19(2), 563–568.

Rintamäki, P. T., Höglund, J., Karvonen, E., Alatalo, R. V., Björklund, N., Lundberg, A., Rätti, O., & Vouti, J. (2000). Combs and sexual selection in black grouse (Tetrao tetrix). Behavioral Ecology, 11(5), 465–471.

Rodríguez-Muñoz, R., Mirol, P. M., Segelbacher, G., Fernández, A., & Tregenza, T. (2007). Genetic differentiation of an endangered capercaillie (Tetrao urogallus) population at the Southern edge of the species range. Conservation Genetics, 8(3), 659–670.

Ronce, O. (2007). How does it feel to be like a rolling stone? Ten questions about dispersal evolution. Annu. Rev. Ecol. Evol. Syst., 38, 231–253.

Rousset, F. (2008). genepop’007: a complete re–implementation of the genepop software for Windows and Linux. Molecular ecology resources, 8(1), 103–106.

Ryder, T. B., Tori, W. P., Blake, J. G., Loiselle, B. A., & Parker, P. G. (2009). Mate choice for genetic quality: a test of the heterozygosity and compatibility hypotheses in a lek-breeding bird. Behavioral Ecology, 21(2), 203–210.

Saccheri, I., Kuussaari, M., Kankare, M., Vikman, P., Fortelius, W., & Hanski, I. (1998). Inbreeding and extinction in a butterfly metapopulation. Nature, 392(6675), 491.

Segelbacher, G., & Storch, I. (2002). Capercaillie in the Alps: genetic evidence of metapopulation structure and population decline. Molecular Ecology, 11(9), 1669–1677.

Segelbacher, G., Höglund, J., & Storch, I. (2003). From connectivity to isolation: genetic consequences of population fragmentation in capercaillie across Europe. Molecular Ecology, 12(7), 1773–1780.

Selwood, K. E., McGeoch, M. A., & Mac Nally, R. (2015). The effects of climate change and landuse change on demographic rates and population viability. Biological Reviews, 90(3), 837-853.

Spielman, D., Brook, B. W., & Frankham, R. (2004). Most species are not driven to extinction before genetic factors impact them. Proceedings of the National Academy of Sciences, 101(42), 15261–15264.

Spottiswoode, C., & Møller, A. P. (2004). Genetic similarity and hatching success in birds. Proceedings of the Royal Society of London B: Biological Sciences, 271(1536), 267–272.

Stephens, P. A., & Sutherland, W. J. (1999). Consequences of the Allee effect for behaviour, ecology and conservation. Trends in ecology & evolution, 14(10), 401–405.

Stephens, P. A., Sutherland, W. J., & Freckleton, R. P. (1999). What is the Allee effect?. Oikos, 185–190.

Stiver, J. R., Apa, A. D., Remington, T. E., & Gibson, R. M. (2008). Polygyny and female breeding failure reduce effective population size in the lekking Gunnison sage-grouse. Biological Conservation, 141(2), 472–481.

Storch I. 2007. Grouse: Status Survey and Conservation Action Plan 2006 –2010. Page 114p. (Storch I, editor). Gland, Switzerland: IUCN and Fordingbridge, UK: World Pheasant Association.

Stubberud, M. W., Myhre, A. M., Holand, H., Kvalnes, T., Ringsby, T. H., Sæther, B. E., & Jensen, H. (2017). Sensitivity analysis of effective population size to demographic parameters in house sparrow populations. Molecular ecology, 26(9), 2449–2465.

Szűcs, M., Melbourne, B. A., Tuff, T., Weiss-Lehman, C., & Hufbauer, R. A. (2017). Genetic and demographic founder effects have long-term fitness consequences for colonising populations. Ecology letters, 20(4), 436–444.

Tanaka, Y. (1997). Extinction of populations due to inbreeding depression with demographic disturbances. Researches on Population Ecology, 39(1), 57–66.

Thomas, C. D., & Kunin, W. E. (1999). The spatial structure of populations. Journal of Animal Ecology, 68(4), 647–657.

Tournier, E., Besnard, A., Tournier, V., & Cayuela, H. (2017). Manipulating waterbody hydroperiod affects movement behaviour and occupancy dynamics in an amphibian. Freshwater Biology, 62(10), 1768–1782.

Tregenza, T., & Wedell, N. (2000). Genetic compatibility, mate choice and patterns of parentage: invited review. Molecular Ecology, 9(8), 1013–1027.

Young, A. G., Clarke, G. M., & Cowlishaw, G. (Eds.). (2000). Genetics, demography and viability of fragmented populations (Vol. 4). Cambridge University Press.

Wang, J., & Whitlock, M. C. (2003). Estimating effective population size and migration rates from genetic samples over space and time. Genetics, 163(1), 429–446.

Waples, R. S., & Yokota, M. (2006). Temporal estimates of effective population size in species with overlapping generations. Genetics.

Wegge P, Larsen BB, Gjerde I, Kastdalen L, Rolstad J & Storaas T. 1987. Natural mortality and predation of adult capercaillie in southeast Norway. Lovel T, Hudson P (eds), 49–56

Westemeier RL, Brawn JD, Simpson SA, Esker TL, Jansen RW, Walk JW, Kershner EL, Bouzat JL, Paige KN. 1998. Tracking the long-term decline and recovery of an isolated population. Science, 282:1695–1698.

Wilson, G. A., & Rannala, B. (2003). Bayesian inference of recent migration rates using multilocus genotypes. Genetics, 163(3), 1177–1191.

Wootton, J. T., & Pfister, C. A. (2013). Experimental separation of genetic and demographic factors on extinction risk in wild populations. Ecology, 94(10), 2117–2123.

