## Supplementary material for "Demography, genetic, and extinction process in a spatially structured population of lekking bird": Sup Mat1

**Supplementary material S1**

**Table 1**. Number of individuals (males and females) captured each year during the 6-years study period (2010-2015).

|  | 2010 | 2011 | 2012 | 2013 | 2014 | 2015 |
| --- | --- | --- | --- | --- | --- | --- |
| Male | 37 | 30 | 28 | 20 | 19 | 24 |
| Female | 29 | 17 | 24 | 17 | 13 | 15 |
