## Supplementary material for "Demography, genetic, and extinction process in a spatially structured population of lekking bird": Sup Mat2

**Supplementary material S2**

**Table S1**. Model selection procedure for survival, dispersal and recapture probability. r = model rank, k = number of model parameters, AICc = Akaike information criterion adjusted for small sample size, ΔAICc = difference of AICc points between the best-supported model and the model considered, w = AICc weight. S = survival probability, E = departure probability, A = arrival probability, P = recapture probability, si = lek size, sex = sex effect, y = year.

| r | Model | k | Deviance | AICc | ΔAICc | w |
| --- | --- | --- | --- | --- | --- | --- |
| 1 | S(si), E(si), A(.), P(sex) | 10 | 1276.46 | 1297.10 | 0.00 | 0.27 |
| 2 | S(.), E(si), A(.), P(sex) | 9 | 1278.79 | 1297.31 | 0.21 | 0.24 |
| 3 | S(.), E(.), A(.), P(sex) | 8 | 1283.04 | 1299.46 | 2.36 | 0.08 |
| 4 | S(si), E(.), A(.), P(sex) | 9 | 1281.10 | 1299.63 | 2.53 | 0.08 |
| 5 | S(si + y), E(si), A(.), P(sex) | 14 | 1271.66 | 1300.90 | 3.80 | 0.04 |
| 6 | S(y), E(si), A(.), P(sex) | 13 | 1273.90 | 1300.97 | 3.87 | 0.04 |
| 7 | S(.), E(si), A(.), P(y + sex) | 15 | 1269.68 | 1301.11 | 4.00 | 0.04 |
| 8 | S(si), E(si), A(.), P(y + sex) | 16 | 1267.66 | 1301.28 | 4.18 | 0.03 |
| 9 | S(si), E(si + sex), A(sex), P(sex) | 12 | 1276.37 | 1301.29 | 4.19 | 0.03 |
| 10 | S(.), E(si + sex), A(sex), P(sex) | 11 | 1278.70 | 1301.47 | 4.37 | 0.03 |
| 11 | S(y), E(.), A(.), P(sex) | 12 | 1278.16 | 1303.08 | 5.97 | 0.01 |
| 12 | S(.), E(.), A(.), P(y + sex) | 14 | 1273.94 | 1303.18 | 6.08 | 0.01 |
| 13 | S(si + y), E(.), A(.), P(sex) | 13 | 1276.28 | 1303.35 | 6.25 | 0.01 |
| 14 | S(.), E(sex), A(sex), P(sex) | 10 | 1282.90 | 1303.55 | 6.44 | 0.01 |
| 15 | S(si), E(.), A(.), P(y + sex) | 15 | 1272.28 | 1303.70 | 6.60 | 0.01 |
| 16 | S(si), E(sex), A(sex), P(sex) | 11 | 1280.98 | 1303.75 | 6.65 | 0.01 |
| 17 | S(y), E(si), A(.), P(y + sex) | 19 | 1263.97 | 1304.25 | 7.15 | 0.01 |
| 18 | S(si + y), E(si), A(.), P(y + sex) | 20 | 1262.15 | 1304.68 | 7.58 | 0.01 |
| 19 | S(si + y), E(si + sex), A(sex), P(sex) | 16 | 1271.57 | 1305.19 | 8.09 | 0.00 |
| 20 | S(y), E(si + sex), A(sex), P(sex) | 15 | 1273.81 | 1305.24 | 8.14 | 0.00 |
| 21 | S(.), E(si + sex), A(sex), P(y + sex) | 17 | 1269.59 | 1305.42 | 8.32 | 0.00 |
| 22 | S(si), E(si + sex), A(sex), P(y + sex) | 18 | 1267.58 | 1305.63 | 8.53 | 0.00 |
| 23 | S(y), E(.), A(.), P(y + sex) | 18 | 1268.23 | 1306.27 | 9.17 | 0.00 |
| 24 | S(si + y), E(.), A(.), P(y + sex) | 19 | 1266.75 | 1307.03 | 9.93 | 0.00 |
| 25 | S(y), E(sex), A(sex), P(sex) | 14 | 1278.02 | 1307.26 | 10.16 | 0.00 |
| 26 | S(.), E(sex), A(sex), P(y + sex) | 16 | 1273.80 | 1307.42 | 10.31 | 0.00 |
| 27 | S(si + y), E(sex), A(sex), P(sex) | 15 | 1276.16 | 1307.58 | 10.48 | 0.00 |
| 28 | S(si), E(sex), A(sex), P(y + sex) | 17 | 1272.16 | 1307.98 | 10.88 | 0.00 |
| 29 | S(y), E(si + sex), A(sex), P(y + sex) | 21 | 1263.88 | 1308.67 | 11.57 | 0.00 |
| 30 | S(si + y), E(si + sex), A(sex), P(y + sex) | 22 | 1262.06 | 1309.13 | 12.03 | 0.00 |
| 31 | S(y), E(sex), A(sex), P(y + sex) | 20 | 1268.08 | 1310.61 | 13.51 | 0.00 |
| 32 | S(si + y), E(sex), A(sex), P(y + sex) | 21 | 1266.62 | 1311.42 | 14.31 | 0.00 |
| 33 | S(.), E(si), A(.), P(.) | 8 | 1296.88 | 1313.30 | 16.20 | 0.00 |
| 34 | S(si), E(si), A(.), P(.) | 9 | 1295.12 | 1313.64 | 16.54 | 0.00 |
| 35 | S(.), E(.), A(.), P(.) | 7 | 1301.14 | 1315.47 | 18.36 | 0.00 |
| 36 | S(si), E(.), A(.), P(.) | 8 | 1299.70 | 1316.12 | 19.02 | 0.00 |
| 37 | S(.), E(si), A(.), P(y) | 13 | 1289.28 | 1316.36 | 19.26 | 0.00 |
| 38 | S(y), E(si), A(.), P(.) | 12 | 1291.67 | 1316.59 | 19.49 | 0.00 |
| 39 | S(si), E(si), A(.), P(y) | 14 | 1287.66 | 1316.90 | 19.80 | 0.00 |
| 40 | S(si + y), E(si), A(.), P(.) | 13 | 1289.98 | 1317.05 | 19.95 | 0.00 |
| 41 | S(.), E(si + sex), A(sex), P(.) | 10 | 1296.80 | 1317.44 | 20.34 | 0.00 |
| 42 | S(si), E(si + sex), A(sex), P(.) | 11 | 1295.03 | 1317.80 | 20.70 | 0.00 |
| 43 | S(.), E(.), A(.), P(y) | 12 | 1293.54 | 1318.46 | 21.36 | 0.00 |
| 44 | S(y), E(.), A(.), P(.) | 11 | 1295.93 | 1318.70 | 21.60 | 0.00 |
| 45 | S(si), E(.), A(.), P(y) | 13 | 1292.21 | 1319.29 | 22.18 | 0.00 |
| 46 | S(si + y), E(.), A(.), P(.) | 12 | 1294.53 | 1319.45 | 22.35 | 0.00 |
| 47 | S(.), E(sex), A(sex), P(.) | 9 | 1301.00 | 1319.52 | 22.42 | 0.00 |
| 48 | S(y), E(si), A(.), P(y) | 17 | 1283.91 | 1319.74 | 22.64 | 0.00 |
| 49 | S(si), E(sex), A(sex), P(.) | 10 | 1299.58 | 1320.22 | 23.12 | 0.00 |
| 50 | S(si + y), E(si), A(.), P(y) | 18 | 1282.41 | 1320.45 | 23.35 | 0.00 |
| 51 | S(.), E(si + sex), A(sex), P(y) | 15 | 1289.20 | 1320.62 | 23.52 | 0.00 |
| 52 | S(y), E(si + sex), A(sex), P(.) | 14 | 1291.58 | 1320.83 | 23.73 | 0.00 |
| 53 | S(si), E(si + sex), A(sex), P(y) | 16 | 1287.57 | 1321.19 | 24.08 | 0.00 |
| 54 | S(si + y), E(si + sex), A(sex), P(.) | 15 | 1289.88 | 1321.30 | 24.20 | 0.00 |
| 55 | S(y), E(.), A(.), P(y) | 16 | 1288.17 | 1321.79 | 24.69 | 0.00 |
| 56 | S(.), E(sex), A(sex), P(y) | 14 | 1293.40 | 1322.64 | 25.54 | 0.00 |
| 57 | S(si + y), E(.), A(.), P(y) | 17 | 1286.95 | 1322.77 | 25.67 | 0.00 |
| 58 | S(y), E(sex), A(sex), P(.) | 13 | 1295.79 | 1322.86 | 25.76 | 0.00 |
| 59 | S(si), E(sex), A(sex), P(y) | 15 | 1292.09 | 1323.51 | 26.41 | 0.00 |
| 60 | S(si + y), E(sex), A(sex), P(.) | 14 | 1294.41 | 1323.65 | 26.55 | 0.00 |
| 61 | S(y), E(si + sex), A(sex), P(y) | 19 | 1283.83 | 1324.11 | 27.01 | 0.00 |
| 62 | S(si + y), E(si + sex), A(sex), P(y) | 20 | 1282.31 | 1324.84 | 27.74 | 0.00 |
| 63 | S(y), E(sex), A(sex), P(y) | 18 | 1288.03 | 1326.08 | 28.97 | 0.00 |
| 64 | S(si + y), E(sex), A(sex), P(y) | 19 | 1286.82 | 1327.10 | 30.00 | 0.00 |

**Table S2**. Model selection procedure for recruitment, dispersal and recapture probability. r = model rank, k = number of model parameters, AICc = Akaike information criterion adjusted for small sample size, ΔAICc = difference of AICc points between the best-supported model and the model considered, w = AICc weight. R = recruitment probability, E = departure probability, A = arrival probability, P = recapture probability, si = lek size, sex = sex effect, y = year.

| r | Model | k | Deviance | AICc | ΔAICc | w |
| --- | --- | --- | --- | --- | --- | --- |
| 1 | R(.), E(si + sex), A(sex), P(sex) | 11 | 1198.98 | 1221.76 | 0.00 | 0.19 |
| 2 | R(.), E(si + sex), A(sex), P(y + sex) | 17 | 1185.94 | 1221.77 | 0.02 | 0.19 |
| 3 | R(y), E(si + sex), A(sex), P(y + sex) | 21 | 1177.83 | 1222.62 | 0.86 | 0.13 |
| 4 | R(si), E(si + sex), A(sex), P(sex) | 12 | 1197.71 | 1222.63 | 0.87 | 0.13 |
| 5 | R(si), E(si + sex), A(sex), P(y + sex) | 18 | 1184.59 | 1222.63 | 0.88 | 0.13 |
| 6 | R(y), E(si + sex), A(sex), P(sex) | 15 | 1191.40 | 1222.83 | 1.07 | 0.11 |
| 7 | R(si + y), E(si + sex), A(sex), P(y + sex) | 22 | 1177.05 | 1224.12 | 2.37 | 0.06 |
| 8 | R(si + y), E(si + sex), A(sex), P(sex) | 16 | 1190.67 | 1224.29 | 2.54 | 0.05 |
| 9 | R(.), E(si + sex), A(sex), P(.) | 10 | 1210.82 | 1231.46 | 9.71 | 0.00 |
| 10 | R(y), E(si + sex), A(sex), P(.) | 14 | 1202.45 | 1231.69 | 9.93 | 0.00 |
| 11 | R(si), E(si + sex), A(sex), P(.) | 11 | 1209.56 | 1232.33 | 10.58 | 0.00 |
| 12 | R(si + y), E(si + sex), A(sex), P(.) | 15 | 1201.78 | 1233.21 | 11.45 | 0.00 |
| 13 | R(y), E(si + sex), A(sex), P(y) | 19 | 1194.83 | 1235.11 | 13.36 | 0.00 |
| 14 | R(.), E(si + sex), A(sex), P(y) | 15 | 1203.94 | 1235.36 | 13.60 | 0.00 |
| 15 | R(si), E(si + sex), A(sex), P(y) | 16 | 1202.67 | 1236.29 | 14.53 | 0.00 |
| 16 | R(si + y), E(si + sex), A(sex), P(y) | 20 | 1194.16 | 1236.69 | 14.94 | 0.00 |

**Table S3**. Model selection procedure for survival and individual inbreeding. r = model rank, k = number of model parameters, AICc = Akaike information criterion adjusted for small sample size, ΔAICc = difference of AICc points between the best-supported model and the model considered, w = AICc weight. S = survival probability, P = recapture probability, PHt = the proportion of heterozygous loci in an individual, IR = the internal relatedness, HL = the homozygosity by locus.

| r | Model | k | Deviance | AICc | ΔAICc | w |
| --- | --- | --- | --- | --- | --- | --- |
| 1 | S(.), P(sex) | 4 | 749.42 | 757.56 | 0.00 | 0.47 |
| 2 | S(PHt), P(sex) | 5 | 749.25 | 759.46 | 1.90 | 0.18 |
| 3 | S(IR), P(sex) | 5 | 749.26 | 759.47 | 1.91 | 0.18 |
| 4 | S(HL), P(sex) | 5 | 749.41 | 759.63 | 2.07 | 0.17 |

**Table S4**. Model selection procedure for recruitment and individual inbreeding. r = model rank, k = number of model parameters, AICc = Akaike information criterion adjusted for small sample size, ΔAICc = difference of AICc points between the best-supported model and the model considered, w = AICc weight. R = recruitment probability, P = recapture probability, PHt = the proportion of heterozygous loci in an individual, IR = the internal relatedness, HL = the homozygosity by locus.

| r | Model | k | Deviance | AICc | ΔAICc | w |
| --- | --- | --- | --- | --- | --- | --- |
| 1 | R(.), P(sex) | 4 | 656.04 | 665.04 | 0.00 | 0.38 |
| 2 | R(IR), P(sex) | 5 | 655.54 | 665.75 | 0.71 | 0.27 |
| 3 | R(PHt), P(sex) | 5 | 656.33 | 666.55 | 1.50 | 0.18 |
| 4 | R(HL), P(sex) | 5 | 656.45 | 666.66 | 1.62 | 0.17 |
