## Supplementary material for "Demography, genetic, and extinction process in a spatially structured population of lekking bird": Sup Mat3

**Supplementary material S3**

In this appendix, we examined the effects on landscape predictors (described below) and genetic divergence among leks. Our main goal was to show that gene flow was non-random in the spatially structured population of *Tetrao urogallus*.

**Preparation of landscape layers and distance matrices**

Extraction and computation of landscape layers were conducted using ArcGIS 10.1 and R (R Core Team 2018). All layers were set at a 25 m resolution.

- The *Slope* layer was computed from a Digital Elevation Model DEM (25m resolution) using the Spatial Analyst toolbox, as in Cayuela et al. (submitted). The layer was rescaled to range from 1 to 100 and we used CIRCUITSCAPE 3.5.8 to compute pairwise effective distances between individual locations, with the layer coded in resistances (higher slope values denoting greater resistance to movement).
- The *Ridge* layer was also computed from the DEM using the Spatial Analyst toolbox. As in Cayuela et al. (*submitted*), we first applied the Focal Statistics tool to compute, for each pixel from the DEM, the mean elevation values within a 1000m neighborhood. We then subtracted the Focal Stats layer from the original DEM and identified ridges as pixels associated with a positive value (pixels above the average elevation of their neighborhood). Elevation values of ridge pixels were rescaled to range from 1(highest elevation) to 100 (lowest elevation) whereas non-ridge pixels were systematically set to 100. We finally used CIRCUITSCAPE 3.5.8 to compute pairwise effective distances between individual locations, with the layer coded in resistances (non-ridge areas denoting greater resistance to movement).
- *Urban* and *Roads* layers were computed from vectorial data (source: BD TOPO® (2013) IGN) as the density of buildings (respectively, of roads) within a 100 m radius buffer. For each layer, densities were rescaled to range from 1 to 100 and we used CIRCUITSCAPE 3.5.8 to compute pairwise effective distances between individual locations, with the layers coded in resistances (higher densities in buildings or roads denoting greater resistance to movement).
- The *NDVI* (Normalized Difference Vegetation Index) was calculated from Landsat 8 images (30 m resolution) taken in August 8th 2016 (source: usgs.gov) using the R package RStoolbox (Leutner, Horning & Schwalb-Willmann, 2018). NDVI values were rescaled to range from 1 to 100 and we used CIRCUITSCAPE 3.5.8 to compute pairwise effective distances between individual locations, with the layer coded in conductances (higher NDVI values denoting forested areas and thus lower resistance to movement).
- The topographic distances *Dtopo* were computed as in Cayuela *et al.* (submitted) as the cumulated length of segments that would be travelled by an individual flying in straight line from a point A to a point B while following topographic relief. From the DEM, we computed the effective travelled distance *d* across each pixel *i* crossed by an individual along its straight line trajectory using the Pythagorean Theorem as follows: $d_{i}=\sqrt{\left( x_{i-1}-x_{i} \right)^{2}+\left( y_{i-1}-y_{i} \right)^{2}}$, with $\left( x_{i-1}-x_{i} \right)=25$ (the length of a pixel), $y_{i-1}$ the altitude at the previous pixel along the trajectory and $y_{i}$the altitude at the focal pixel. For each pair of points, the topographic distance *Dtopo* was then computed as the sum of distances $d_{i}$along the trajectory.
- Euclidean distances *ED* were finally computed in CIRCUITSCAPE 3.5.8 from a null raster, with all pixels set to 1 and the whole layer coded in resistances.

**Statistical analyses and results**

Following Cayuela *et al.* (submitted), all the genetic analyses were performed separately on males (dataset M) and females (dataset F) with the Bray-Curtis percentage dissimilarity metric (Bc; Legendre & Legendre, 1998) as an inter-individual measure of pairwise genetic distances. For each sex, we designed a complete model as follows: $GD \sim ED+Dtopo+Slope+Ridge+Urban+Roads+NDVI.$ We did not run this model on all possible pairs of individuals, but selected a subset of pairwise data corresponding to individuals located less than 14 and 23 km apart, in males and females, respectively. These distances were previously found to optimize the amount of variance explained in measures of genetic differentiation in each sex (Cayuela et al., submitted).

We coupled multiple linear regressions on standardized data with commonality analyses (CA; (Ray-Mukherjee *et al.*, 2014) to identify the main contributors to the variance in the dependent variables after sequential removing of suppressors and unnecessary predictors. Predictors were identified as suppressors when their unique contribution was (almost) totally counterbalanced by a negative commonality coefficient (classical suppression) or when standardized regression coefficients and structure coefficients were of opposite signs (cross-over suppression; Paulhus *et al.*, 2004; Prunier *et al.*, 2017). Predictors were identified as unnecessary when their unique contribution (U) was null (or when the lower bound of 95 % confidence intervals CI around U was null), indicating that they only contributed to the variance in the dependent variable because of their synergistic association with one or several other predictors (Prunier *et al.*, 2015). The 95 % CI around beta coefficients β and unique contributions U were computed from bootstrap resampling (1000 iterations).

For each sex (M and F), the following table provides details about runs of identification of unnecessary predictors (in synergistic association with other predictors) and suppressors in full models. Are provided: typical results of the different runs of multiple linear regressions (model fit R², predictors Pred, structure coefficients rs and standardized coefficents β), along with additional parameters derived from CA: unique, common and total contributions of predictors to the variance in dependent variable (U, C and T). The table also provides 95% confidence intervals about standardized coefficents β (as β–CI_low_ and β–CI_up_) and about U (U–CI_low_ and U–CI_up_) as computed from bootstrap (1000 iterations). The rationale for withdrawal of predictors (Ra) is the following: S: synergistic association with other predictors (U–CI_low_ = 0); CO: Cross-over suppression; CL: Classical suppression. In italics: parameters allowing the identification of unnecessary predictors and suppressors. In bold: main contributors to the variance in measures of genetic distances.

Despite the addition of several predictors likely to affect patterns of genetic differentiation in *Tetrao urogallus* (*NDVI*, *Roads* and *Urban*), we found the exact same results as in Cayuela et al. (*submitted*). Whatever the sex, the main contributor to the variance in measures of genetic differentiation was the topographic distance. In males, genetic distances were also affected by slopes. Anthropogenic features likely to alter *T. urogallus* habitat were not retained as important predictors of genetic structure in this species. The amount of explained variance was much higher in males (R² = 0.305) than in females (R² = 0.031), confirming a strong spatial structure of relatedness in males.

| Sex | Run | R² | Pred | rs | β | β-CI_low_ | β-CI_up_ | U | U-CI_low_ | U-CI_up_ | C | T | Ra |
| --- | --- | --- | --- | --- | --- | --- | --- | --- | --- | --- | --- | --- | --- |
| M | 1 | 0.323 | ED | *0.819* | *-1.010* | -2.159 | 0.201 | 0.005 | 0.000 | 0.021 | 0.211 | 0.216 | CO |
|  |  |  | Slope | 0.824 | 0.303 | 0.184 | 0.416 | 0.033 | 0.012 | 0.060 | 0.186 | 0.219 |  |
|  |  |  | Ridge | 0.800 | 0.203 | -0.017 | 0.406 | 0.005 | 0.000 | 0.019 | 0.202 | 0.207 |  |
|  |  |  | Topo | 0.894 | 0.441 | 0.310 | 0.564 | 0.062 | 0.029 | 0.101 | 0.196 | 0.258 |  |
|  |  |  | NDVI | 0.829 | 1.290 | 0.145 | 2.417 | 0.008 | 0.000 | 0.027 | 0.214 | 0.222 |  |
|  |  |  | Urban | *0.816* | *-0.599* | -1.212 | 0.010 | 0.005 | 0.000 | 0.019 | 0.210 | 0.215 | CO |
|  |  |  | Roads | 0.629 | 0.012 | -0.082 | 0.104 | 0.000 | 0.000 | 0.006 | 0.128 | 0.128 |  |
|  | 2 | 0.309 | Slope | 0.842 | 0.309 | 0.204 | 0.426 | 0.035 | 0.015 | 0.063 | 0.184 | 0.219 |  |
|  |  |  | Ridge | 0.818 | 0.158 | -0.058 | 0.362 | 0.003 | 0.000 | 0.015 | 0.204 | 0.207 |  |
|  |  |  | Topo | 0.914 | 0.394 | 0.271 | 0.531 | 0.055 | 0.026 | 0.099 | 0.203 | 0.258 |  |
|  |  |  | NDVI | *0.848* | *-0.228* | -0.501 | 0.042 | 0.004 | 0.000 | 0.020 | 0.218 | 0.222 | CO |
|  |  |  | Roads | 0.643 | 0.000 | -0.086 | 0.096 | 0.000 | 0.000 | 0.005 | 0.128 | 0.128 |  |
|  | 3 | 0.305 | Slope | 0.847 | 0.263 | 0.160 | 0.364 | 0.031 | 0.011 | 0.057 | 0.188 | 0.219 |  |
|  |  |  | Ridge | 0.823 | 0.012 | -0.117 | 0.131 | 0.000 | 0.000 | 0.006 | 0.207 | 0.207 |  |
|  |  |  | Topo | 0.920 | 0.353 | 0.243 | 0.464 | 0.052 | 0.025 | 0.091 | 0.206 | 0.258 |  |
|  |  |  | Roads | *0.648* | *-0.009* | -0.101 | 0.079 | 0.000 | 0.000 | 0.006 | 0.128 | 0.128 | CO |
|  | 4 | 0.305 | **Slope** | **0.847** | **0.260** | **0.168** | **0.356** | **0.035** | **0.015** | **0.066** | **0.184** | **0.219** |  |
|  |  |  | Ridge | 0.823 | 0.010 | -0.135 | 0.135 | 0.000 | 0.000 | 0.007 | 0.207 | 0.207 |  |
|  |  |  | **Topo** | **0.920** | **0.352** | **0.241** | **0.464** | **0.052** | **0.025** | **0.090** | **0.206** | **0.258** |  |
| F | 1 | 0.073 | ED | 0.402 | 1.989 | 0.572 | 3.345 | 0.012 | 0.001 | 0.034 | 0.000 | 0.012 |  |
|  |  |  | Slope | 0.324 | 0.113 | -0.043 | 0.256 | 0.003 | 0.000 | 0.014 | 0.005 | 0.008 |  |
|  |  |  | Ridge | *0.253* | *-0.305* | -0.540 | -0.085 | 0.010 | 0.001 | 0.029 | -0.005 | 0.005 | CO |
|  |  |  | Topo | 0.648 | 0.303 | 0.167 | 0.423 | 0.031 | 0.009 | 0.059 | 0.000 | 0.031 |  |
|  |  |  | NDVI | *0.379* | *-1.671* | -2.849 | -0.492 | 0.010 | 0.001 | 0.029 | 0.001 | 0.010 | CO |
|  |  |  | Urban | *0.387* | *-0.182* | -1.069 | 0.749 | 0.000 | 0.000 | 0.008 | 0.011 | 0.011 | CO |
|  |  |  | Roads | -0.125 | -0.141 | -0.242 | -0.034 | *0.010* | 0.001 | 0.031 | *-0.009* | 0.001 | CL |
|  | 2 | 0.034 | ED | *0.590* | *-0.137* | -0.325 | 0.061 | 0.003 | 0.000 | 0.018 | 0.009 | 0.012 | CO |
|  |  |  | Slope | 0.475 | 0.060 | -0.081 | 0.212 | 0.001 | 0.000 | 0.013 | 0.007 | 0.008 |  |
|  |  |  | Topo | 0.950 | 0.249 | 0.124 | 0.378 | 0.022 | 0.005 | 0.051 | 0.009 | 0.031 |  |
|  | 3 | 0.031 | Slope | *0.499* | *-0.018* | -0.110 | 0.077 | 0.000 | 0.000 | 0.009 | 0.007 | 0.008 | CO |
|  |  |  | Topo | 0.996 | 0.185 | 0.089 | 0.281 | 0.023 | 0.005 | 0.053 | 0.007 | 0.031 |  |
|  | 4 | 0.031 | **Dtopo** | **0.175** | **0.175** | **0.108** | **0.259** | / | / | / | / | / |  |

REFERENCES

Cayuela*, H., Boualit*, L., Laporte, M., Prunier, J., Foletti, F., Clobert, J., Jacob, G. Kin-dependent dispersal influences relatedness and genetic structuring in a lek system. Submitted to *Journal of Animal Ecology*, submitted.

Legendre P. & Legendre L.F.J. (1998) *Numerical Ecology*, 2nd English ed. Elsevier Science B.V, Amsterdam, Amsterdam.

Leutner B., Horning N. & Schwalb-Willmann J. (2018) *RStoolbox: Tools for Remote Sensing Data Analysis. R package version 0.2.1.*

Paulhus D.L., Robins R.W., Trzesniewski K.H. & Tracy J.L. (2004) Two replicable suppressor situations in personality research. *Multivariate Behavioral Research* **39**, 303–328.

Prunier J.G., Colyn M., Legendre X. & Flamand M.-C. (2017) Regression commonality analyses on hierarchical genetic distances. *Ecography* **40**, 001–014.

Prunier J.G., Colyn M., Legendre X., Nimon K.F. & Flamand M.C. (2015) Multicollinearity in spatial genetics: separating the wheat from the chaff using commonality analyses. *Molecular Ecology* **24**, 263–283.

Ray-Mukherjee J., Nimon K., Mukherjee S., Morris D.W., Slotow R. & Hamer M. (2014) Using commonality analysis in multiple regressions: a tool to decompose regression effects in the face of multicollinearity. *Methods in Ecology and Evolution* **5**, 320–328.
